# DNA Content and Ploidy Estimation of *Festuca ovina* Accessions by Flow Cytometry

**DOI:** 10.1101/2020.02.06.938100

**Authors:** Yinjie Qiu, Sierra Hamernick, Joan Barreto Ortiz, Eric Watkins

## Abstract

*Festuca ovina* is a fine fescue that is used as a low-input turfgrass. The ploidy levels of *F. ovina* accessions held by the USDA National Plant Germplasm System (NPGS) are unknown, limiting the use of the germplasm in breeding programs. The objective of this study was to determine DNA content and estimate ploidy of these 127 accessions. Among the accessions, we identified a wide range of ploidy levels from diploid to octoploid. We also found the accessions with higher ploidy levels usually had larger seed size. These results will be informative to plant breeders and researchers using germplasm from the *F. ovina* collection and point to challenges in maintaining polyploid, outcrossing germplasm seed stocks in common nurseries.

## INTRODUCTION

Fine fescues (*Festuca* spp. L.) are a diverse group of grasses characterized by fine leaf texture. Fine fescues are native to Eurasia but have been introduced and naturalized to many temperate regions in the world (Barkworth, Capels, Long, Anderton, & Piep, 2007; Beard, 1972; Vasey, 1883). Theses grasses are used for forage, ornamental purposes, and particularly as low-input turfgrasses. The group comprises genetically diverse taxa, including hard fescue (*Festuca brevipila* Tracey, 2n=6x=42), sheep fescue (*F. ovina* L., 2n=4x=28), strong creeping red fescue (*Festuca rubra* ssp. *rubra* 2n=8x=56), slender creeping red fescue [*Festuca rubra* ssp. *litoralis* (G. Mey.) Auquier 2n=6x=42], and Chewings fescue [*Festuca rubra* ssp. *fallax* (Thuill.) Nyman 2n=6x=42] ((Ruemmele, Brilman, & Huff, 1995).

Based on their morphology, cytology, and chloroplast-genome-based phylogenetic relationship, fine fescues are divided in two complexes: the *F. ovina* complex, which includes *F. brevipila* and *F. ovina*, and the *F. rubra* complex, which includes *F. rubra* ssp. *litoralis, F. rubra* ssp. *rubra*, and *F. rubra* ssp. *fallax* (Huff & Palazzo, 1998; Qiu, Hirsch, Yang, & Watkins, 2019; Wilkinson & Stace, 1991). Species in *F. ovina* are bunch-type and non-rhizomatous while subspecies in the *F. rubra* complex can be rhizomatous (ssp. *litoralis* and ssp. *rubra*) and bunch-type (ssp. *fallax*). Taxon identification within the *F. ovina* complex is difficult because of morphological and ecotype diversity (Piper, 1906; Schmit, Duell, & Funk, 1974). Previously, leaf color has been used for species identification; however, in the United States, sheep fescue is described as having a bluish-gray leaf color and hard fescue leaf blade color is considered green (Beard, 1972), while in Europe, it is the opposite (Hubbard, 1968). The *F. ovina* complex also includes some other taxa beyond those commonly used as turfgrasses that have ploidy level varying from diploid (2n=2x=14) such as *Festuca ovina* ssp. *ovina*, tetraploid *Festuca armoricana*, and hexaploid *Festuca huonii* (Seal, 1983; Stace, 2010; Wilkinson & Stace, 1991). All of these factors make the identification of *F. ovina* challenging.

Turfgrass breeding and genetics objectives are focused on aesthetic beauty, disease resistance, drought tolerance, and traits associated with reduced inputs (Bonos, Clarke, & Meyer, 2006; Bonos & Huff, 2013; Casler, 2003; Clarke et al., 2006). Plant breeders seek desirable alleles in exotic accessions and attempt to introgress them into the existing cultivars. One of the most valuable resources for breeders is the USDA-ARS National Plant Germplasm System (NPGS), which holds more than 500,000 accessions that represent more than 10,000 plant species. These accessions have been widely used in plant breeding programs for abiotic and biotic stress improvement (Chang & Hartman, 2017; Christensen et al., 2007; Dilday, Lin, & Yan, 1994; Leng, Wang, Ali, Zhao, & Zhong, 2016; Nelson, Amdor, & Orf, 1987). To improve turfgrass cultivars by utilizing the germplasm accessions, it is important to know the ploidy level of the accessions used to avoid hybridizing plants with different ploidy levels that results in nonviable offspring. Additionally, without knowing the ploidy level, genetics and phenotyping of these germplasm could lead to incorrect interpretation of results.

Traditional plant breeding methods that emphasize hybridizing elite germplasm usually result in the loss of genetic diversity and heterosis which could result in greater susceptibility to important stresses (Christiansen, Andersen, & Ortiz, 2002; Melchinger, 1999; Reif et al., 2005). The use of exotic germplasm has been a common practice in maintaining plant genetic diversity in the breeding process to reduce these problems and avoid breeding bottlenecks (Goodman, 1999; Mikel, Diers, Nelson, & Smith, 2010; Prasanna, 2012; Van Esbroeck & Bowman, 1998). In both cool-season and warm-season turfgrasses, genetic diversity of available germplasm has been studied using either molecular markers or sequencing arrays (Baird et al., 2012; Budak, Shearman, Gaussoin, & Dweikat, 2004; Chen, Wang, Waltz, & Raymer, 2009).

For NPGS accessions from the *F. ovina* complex, it is necessary to determine ploidy level before conducting germplasm selection and performing hybridization; this has traditionally been done by counting chromosomes (Maluszynska, 2003; Vargas, McAllister, Morton, Jury, & Wilkinson, 1999), a reliable but time-consuming method which is made all the more challenging when dealing with the numerous and small (even under magnification) chromosomes of the fine fescue species. Flow cytometry is a powerful tool for DNA content measurement and allows researchers to estimate the ploidy level by comparing to known standards. This method is less time consuming, cheaper, and proven to work in grasses where it has been used to calculate the DNA content and estimate ploidy levels in Texas bluegrass (*Poa arachnifera* Torr.), buffalograss [*Bouteloua dactyloides* (Nutt.) Engelm.], perennial ryegrass (*Lolium perenne* L.) and fine fescues (Ka Arumuganathan & Earle, 1991; Goldman, 2015; Huff & Palazzo, 1998; Johnson, Kenworthy, Auld, & Riordan, 2001; Johnson, Riordan, & Arumuganathan, 1998; Qiu et al., 2019).

We used flow cytometry to determine the DNA content and ploidy level of 127 USDA *F. ovina* PI collections. In addition, we used image analysis to measure and compare the seed size on selected PI accessions among ploidy levels.

## MATERIAL AND METHODS

### Plant Material

A total of 127 accessions labeled as *Festuca ovina* from 20 countries were obtained from the USDA Germplasm Resources Information Network (GRIN) in 2016 (**Table S1**).

Seeds of each accession were sown into greenhouse pots (four-inch size) filled with BRK Promix soil (Premier Tech, USA) at the Plant Growth Facility at the University of Minnesota in St. Paul. After reaching four to five leaf stage, five seedlings per accession were randomly selected and transplanted into individual one-inch size cone container. Plants were grown with 16 h day and 8 hr night with bi-daily watering and weekly fertilization using Peat-lite 20-10-20 fertilizer (J.R. Peters Inc.) with supplemental ammonium sulfate and Sprint 330 (BASF, USA). The five genotypes of each accession were vegetatively cloned into six replications of each genotype and transplanted to a field nursery in at the Minnesota Agricultural Experiment Station in St. Paul, MN. *F. ovina* cv. Quatro and *F. brevipila* cv. Beacon were also planted in the field as standards.

### Flow Cytometry Procedure

To determine the nucleus DNA content of accessions, flow cytometry was carried out using the method described by Arumuganathan and Earle (1991). Because seeds used in this study resulted from open pollination, we evaluated three genotypes for each accession to obtain a more accurate ploidy representation of the population. When the three selected genotypes were not at the same ploidy level, a fourth genotype was evaluated. The ploidy level of the accession was determined by the ploidy of the majority genotypes (75%).

Fresh mature *F. ovina* leaf samples in the nursery field were harvested between 9-11 a.m. and trimmed to 1-2 cm and used for flow cytometry. Perennial ryegrass leaf tissue was harvested at the same time in the greenhouse. For each sample the flow cytometry staining solution contained 4.29 μL propidium iodide, 0.71 mL of CyStain UV Precise P staining buffer, and 2.14 μL RNAseA. To prepare plant tissue, a 0.5 cm × 0.5 cm leaf samples was excised into small pieces using a razor blade in 500 μL CyStain UV Precise P extraction buffer (Sysmex) and passed through a 50-μm size filter (Sysmex). The staining solution was added to the flow-through to stain the nuclei in each sample. Samples were stored on ice before loading the flow cytometer. Flow cytometry was carried out using the BD LSRII H4760 (LSRII) instrument (BD Biosciences, USA) with PI laser detector using 480V with a minimum of 1,000 events at the University of Minnesota Flow Cytometry Resource (UCRF). Data were visualized and analyzed on BD FACSDiva 8.0.1 software.

### DNA Content Estimation and Ploidy Level Determination

Species DNA content was estimated following the method described by Dolezel and Bartoš (2005). The perennial ryegrass (*Lolium perenne*) 2C DNA content (2C = 5.66 pg/2C) served as the diploid DNA content standard (Arumuganathan, Tallury, Fraser, Bruneau, & Qu, 1999). Sample 2C DNA content was calculated using **Equation 1**. To estimate the ploidy level in *F. ovina* complex, diploid *F. ovina* PI 230246 measured in previous study was used (2C = 4.7 pg/2C); species ploidy level was estimated using **Equation 2**.

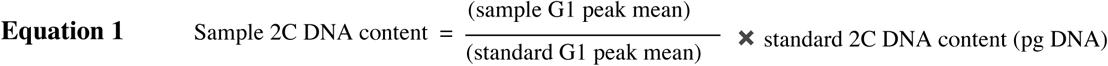

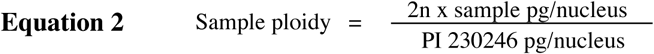

### Seed Size Measurement and Comparison

For seed size measurements, we randomly picked 10 seeds from each of three accessions for each calculated ploidy level (a total of 12 accessions). Tetraploid *F. Ovina* cv. Quatro and hexaploid *F. brevipila* cv. Beacon were also included as references.

Seeds were spread on a digital scanner (Epson perfection v6 flatbed scanner, Nagano, Japan) and scanned at 1200 dots per inch (dpi) with the size of 2097 × 1624 pixels. The images were processed with a custom Matlab script (https://github.umn.edu/jbarreto/seed_morphology) that transformed the original images to measure the seed length and width respectively as the length (in pixels) of the major and minor axes of a fitted ellipse, whereas the area was calculated as the number of pixels in the seed. The seed area was calculated for each of 10 seeds for each of the 14 entries. The seed area was used to analyze the correlation between ploidy level and seed size and visualized using the ggplot2 package in R (Kahle & Wickham, 2013).

## RESULTS

### DNA Content Measurement and Ploidy Estimation

*Festuca ovina* accessions had a high level of DNA content variation with the smallest DNA content of 3.77 pg (2C) from PI 115358, and the largest genome 19.66 pg (2C) from PI 302899 (**Table S2**). Flow cytometry revealed the 127 accessions represented a range of ploidy levels from diploid to octoploid (**Figure 1**).

**Figure 1.**
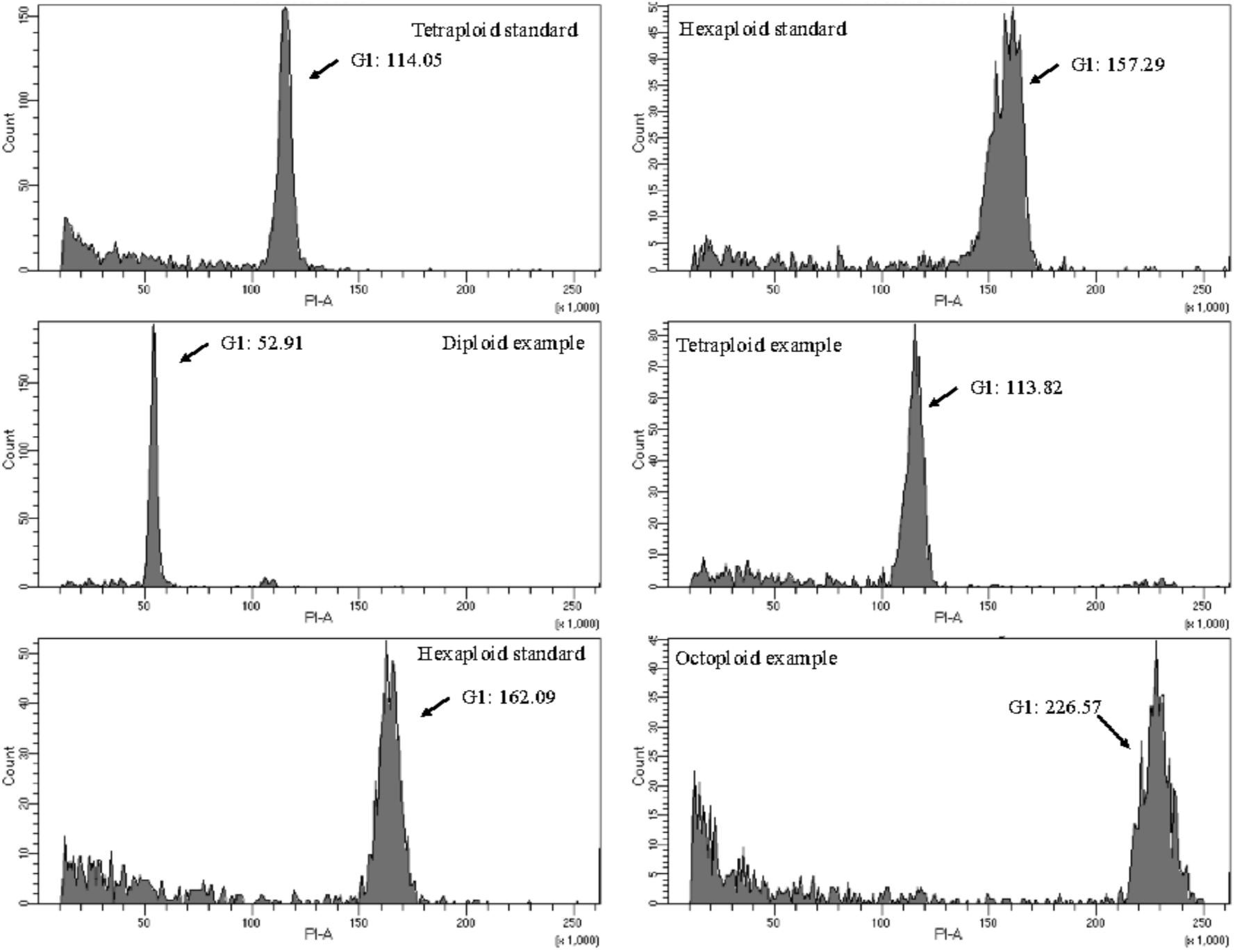
Flow cytometry data example of the PI accessions. Tetraploid sheep fescue cultivar ‘Quatro’ and hexaploid hard fescue cultivar ‘Beacon’ were included as standards. PI accessions included taxa that cover at least four ploidy levels.

The largest standard deviation within accession was 1.25 pg (PI 274619) while the lowest was 0.01 pg (W6 23622). The average of standard deviation of the 126 accessions was 0.45 pg. The majority of accessions had consistent DNA content (judged by standard deviation) within the three genotypes examined. There were cases in which the sampled genotypes had more than one ploidy level. For example, for PI 330706, there were two tetraploids and one triploid plant; however, the fourth sample suggested the accession was tetraploid (**Figure 2**). Three out of the five genotypes examined in PI 235072 accession had three different ploidies (data not shown) and were therefore excluded from further analysis. The DNA content estimation of the 126 accessions is summarized in **Figure 3**.

**Figure 2.**
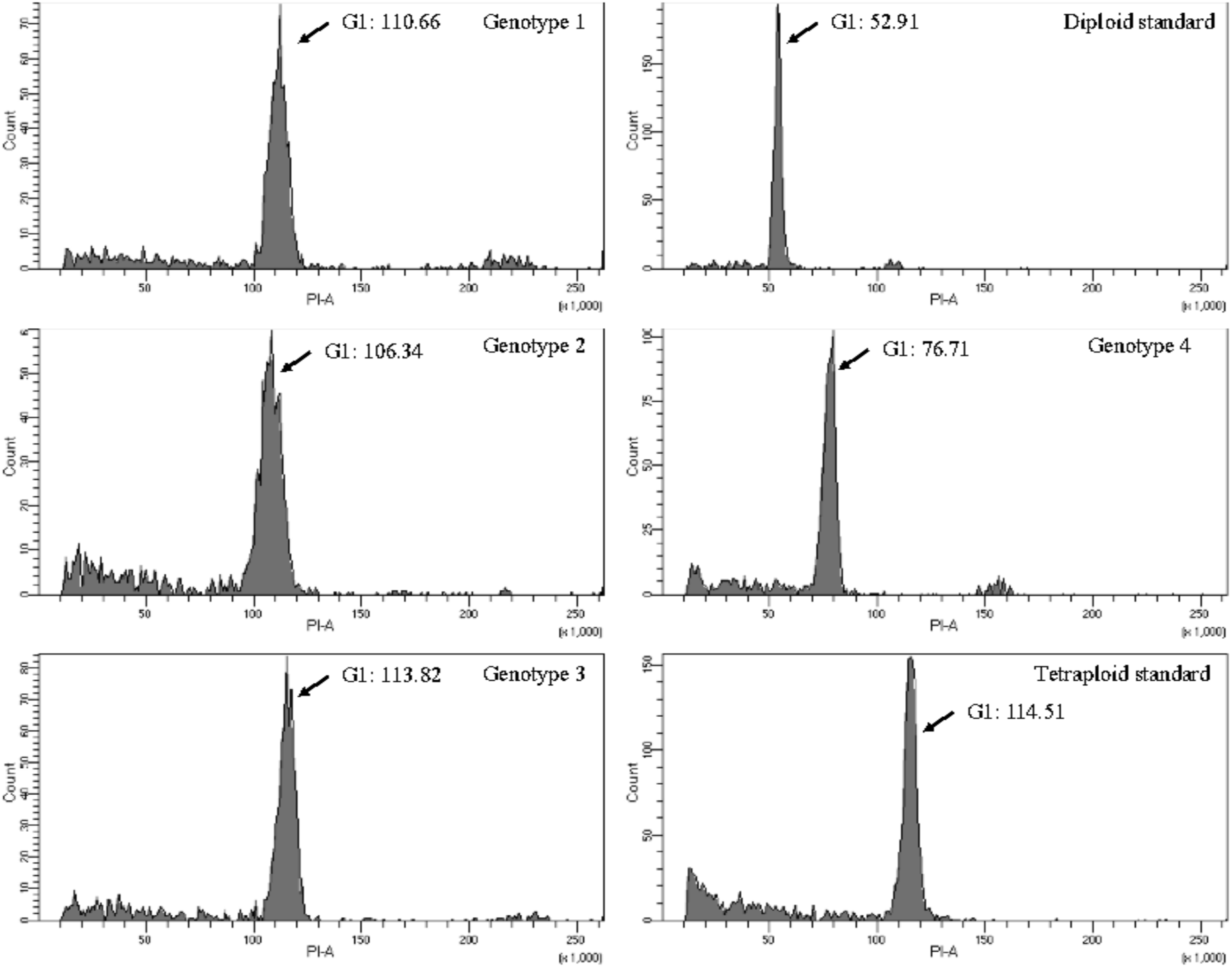
Flow cytometry histogram of PI 330706. Three genotypes from this accession had similar DNA content compared to the tetraploid standard cultivar ‘Quatro’. One genotype from this accession had a DNA content between diploid and tetraploid standard and was estimated to be triploid based on the DNA content estimation.

**Figure 3.**
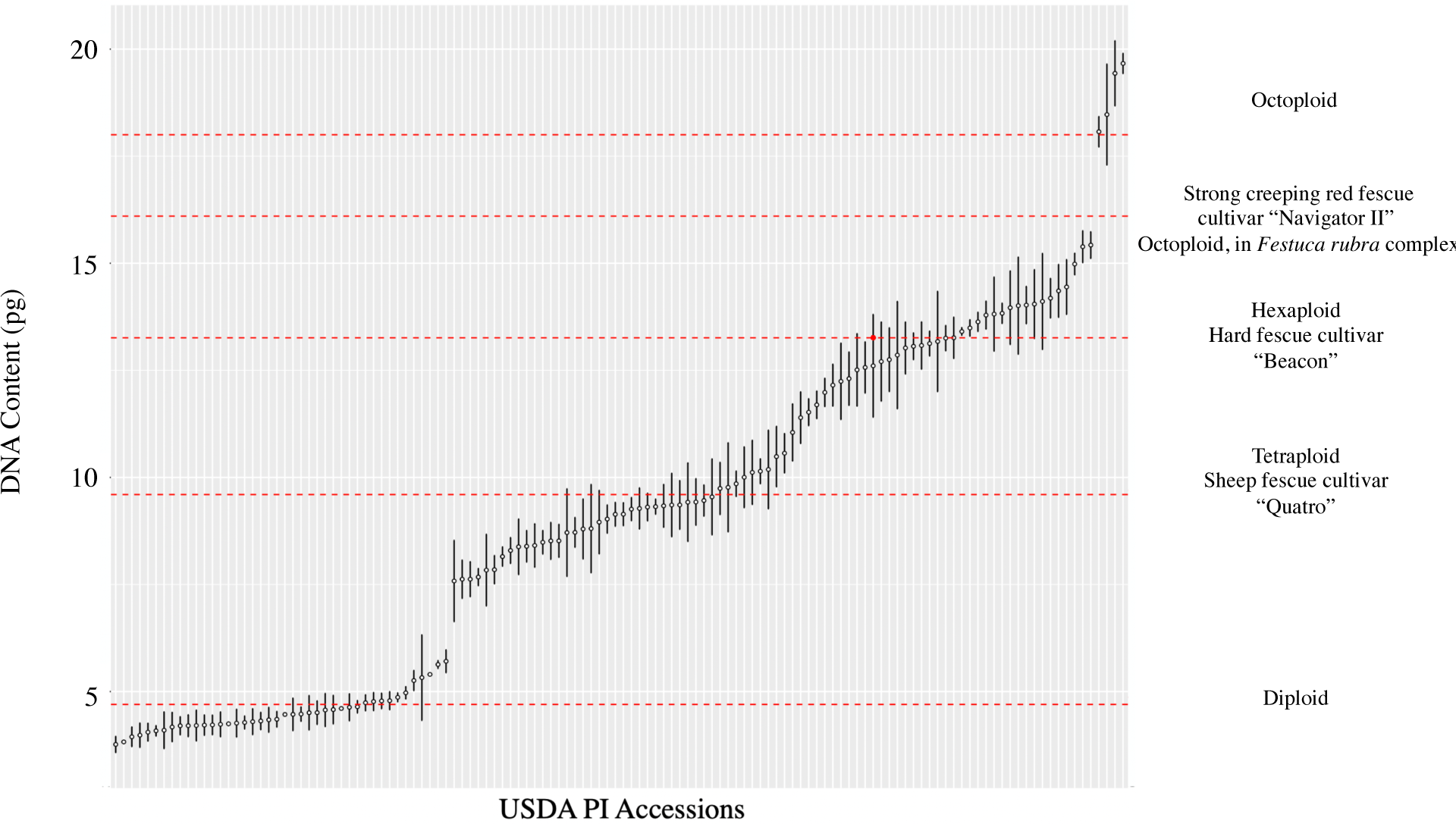
Accessions sorted by the estimated DNA content. Dashed red lines represent the ploidy level estimation for cultivars listed to the right of the figure.

### Ploidy Estimation

To estimate ploidy, estimated DNA content was divided by the DNA content of the known diploid and round up to an integer. Ploidy estimates for the 126 accessions are summarized in **Table 1**. The majority of the accessions (82%) were di-, tetra-, hexa-, and octoploids while 18% of accessions were potential tri-, penta-, and septaploids. The seven accessions estimated as triploid had the estimated ploidy level between 3.23-3.47; the 14 pentaploid accessions had estimated ploidy between 4.70-5.47; and the 2 septaploids had estimated ploidy level of 6.55 and 6.56. Ploidy distribution by country of origin is shown in **Figure 4**.

**Table 1.** The ploidy level estimation of the 126 USDA PI accessions.

| Ploidy level | Diploid | Tetraploid | Hexaploid | Octoploid | Other Ploidy |
| --- | --- | --- | --- | --- | --- |
| Accession Count | 42 | 34 | 22 | 4 | 24 |
| DNA content range (pg) | 3.76 ~ 5.71 | 8.29 ~ 10.48 | 13.02~ 14.98 | 18.07 ~ 19.66 | NA |

**Figure 4.**
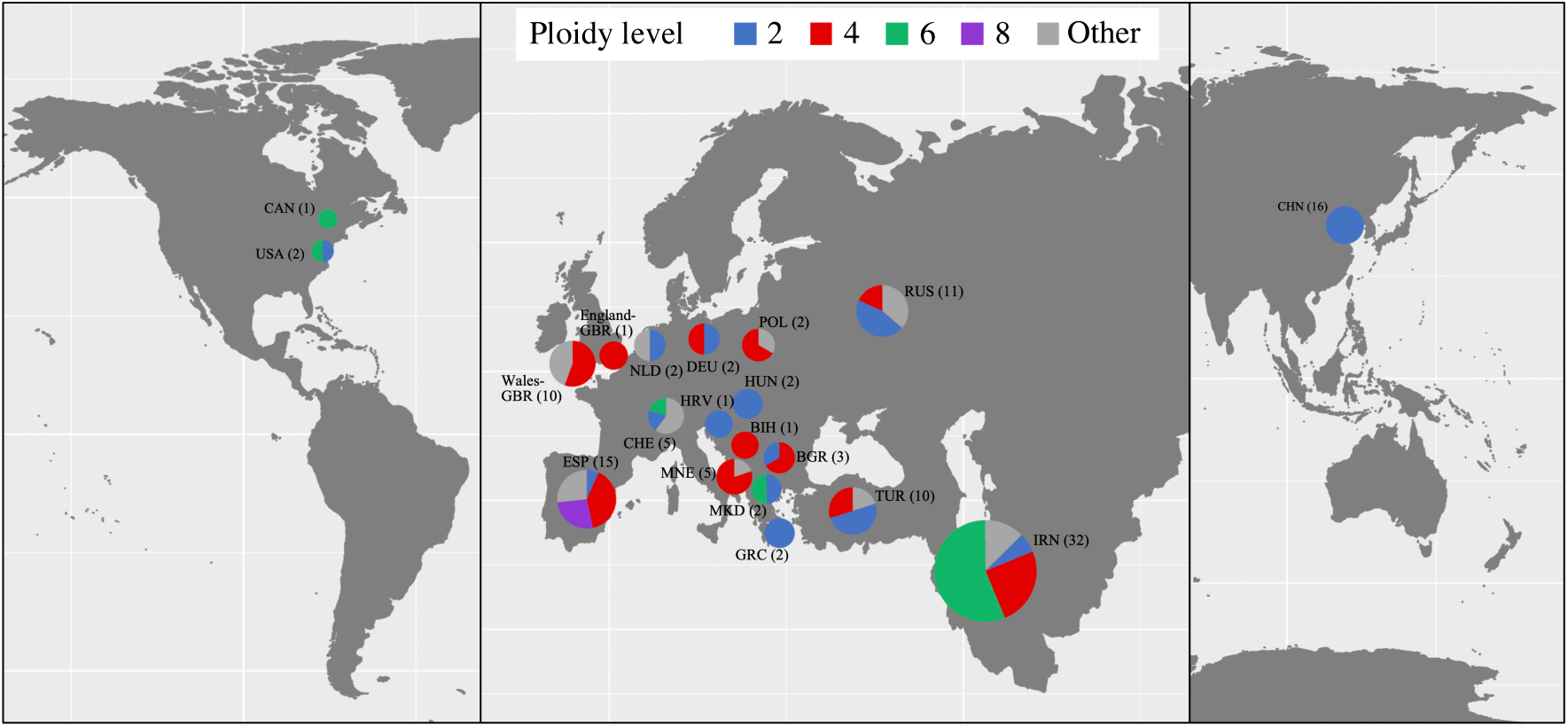
Ploidy estimation by DNA content for the 126 USDA PI accessions.

Besides DNA content variation, we observed leaf color variation at different ploidy levels. For all ploidy levels, there were the presence of blue-greenish color and green color (**Figure 5**).

**Figure 5.**
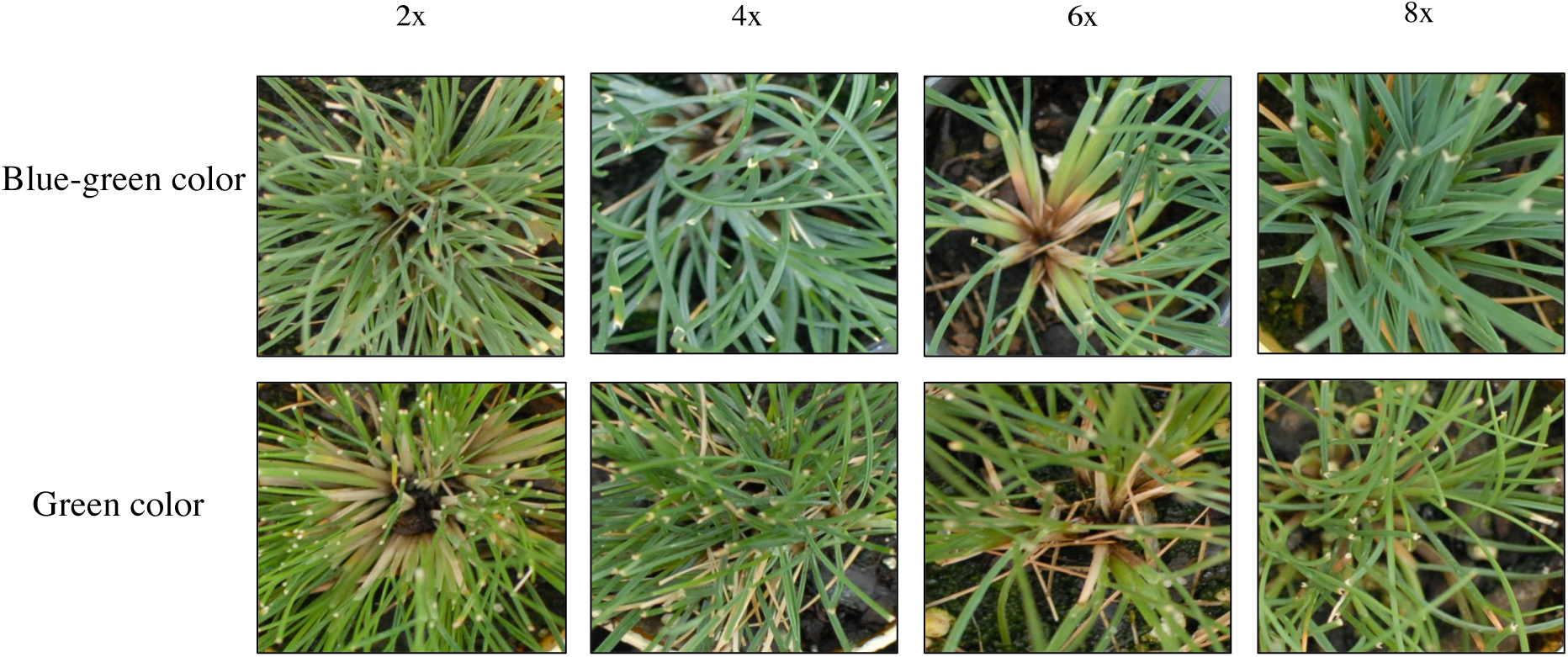
Leaf color differences observed in the *F. ovina* collection. We observed both green and blue-greenish leaf color for plants at all four ploidy levels.

### Seed Size and Ploidy

We found statistical differences in seed size among ploidy levels. In general, higher ploidy level was associated with larger seed size. Diploid accessions had the average seed size between 1.5 and 2.5 mm2, tetraploid accessions seed size varies between 2 and 4 mm2, and hexaploid and octoploid had similar seed size between 4-6.5 mm2 (**Figure 6**).

**Figure 6.**
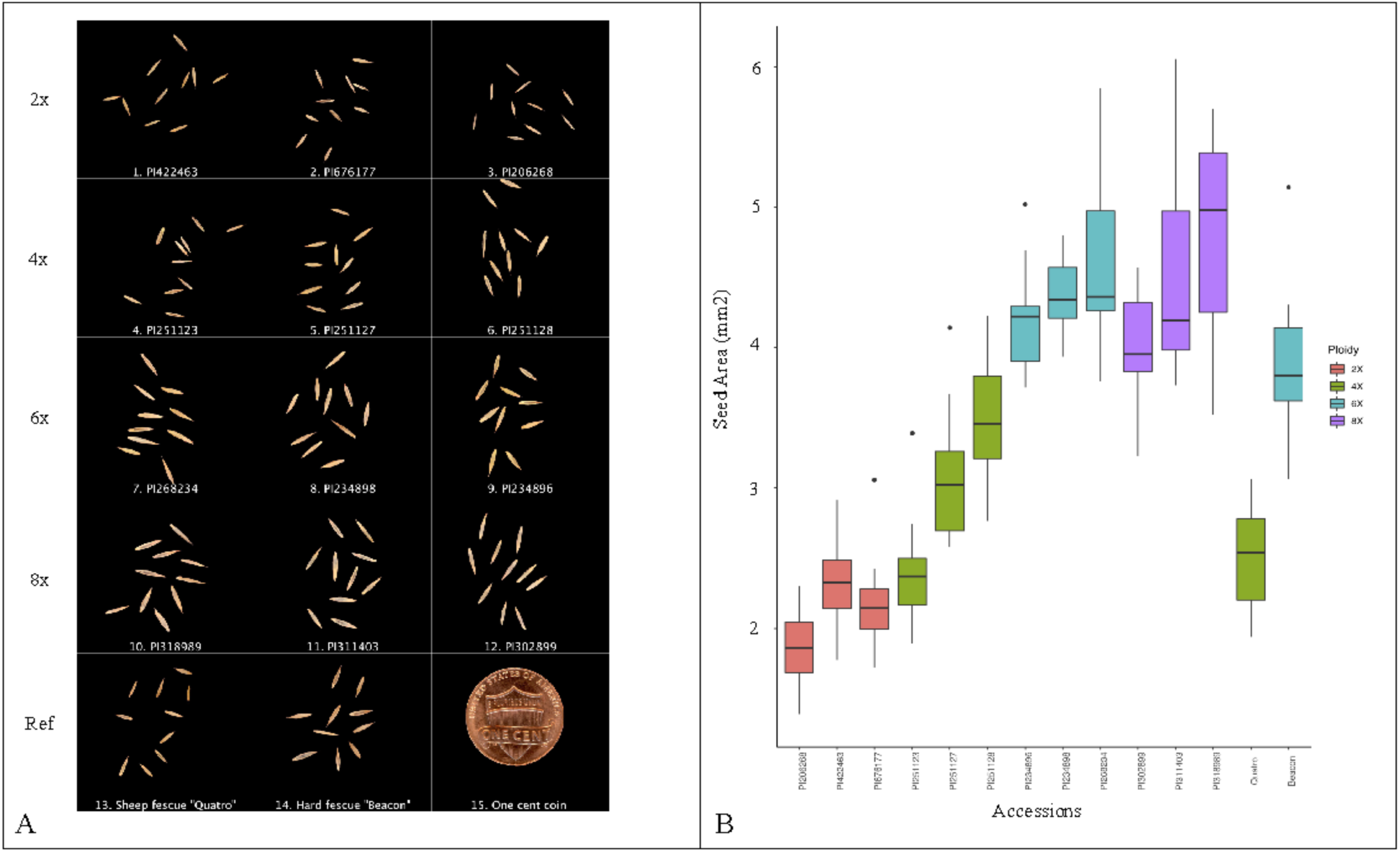
Comparison of seed area and ploidy level. The seed morphology of the examined 12 accessions and two fine fescues cultivars were shown in (A) with the ploidy of each accession labeled on the left. Box plot of seed size comparison is shown in (B); larger seed size was associated with higher ploidy level.

Analysis of the main effects model suggested that the ploidy level explains over 70% of the variation in seed area and length, and more than 55% of the variation in seed length. We rejected the hypothesis that ploidy level has no significant effect on the seed size of fine fescues (*p-value*: < 2.2e-16) (**Table S3, S4)**.

## DISCUSSION

Accessions from the USDA NPGS can serve a valuable role as a source of genetic diversity for crop improvement providing an important gene pool for the improvement of both abiotic and biotic stress tolerance (Rubenstein, Smale, & Widrlechner, 2006). Numerous studies have been done to characterize PI germplasm with topics varying from soybean maturity groups (Nelson et al., 1987) to the identification of allelopathic accessions in rice (Dilday et al., 1994). These accessions have also been used to selected for reproductive characteristics and seed production in garlic (Jenderek & Hannan, 2004), screening for disease resistance in dry beans (Pastor-Corrales, 2003), and the evaluation of drought tolerance in watermelon at seedling stages (Zhang et al., 2011). In forage crops and turfgrasses, tall wheatgrass was evaluated for forage yield and quality (Vogel & Moore, 1998) and Kentucky bluegrass was studied for the reproductive mode (Wieners, Fei, & Johnson, 2006). It is clear that USDA PI accessions provide a diverse gene pool for cultivar abiotic and biotic stress tolerance improvement (Rubenstein et al., 2006). The USDA accessions are particularly important for turfgrass breeding and genetics because commercial turfgrass cultivars have undergone heavy selection and lost genetic diversity.

Different from major crops, most turfgrass species have numerous ploidy levels and some are morphologically indistinguishable; therefore, germplasm characterization is very important. Ploidy determination of 200 perennial ryegrass (*Lolium perenne*) suggested that six accessions were tetraploid and 194 were diploid (Wang, Bigelow, & Jiang, 2009). Screening of buffelgrass (*Cenchrus ciliaris* L. syn. *Pennisetum* ciliare (L.) Link) germplasm suggested multiple ploidy levels existed in the PI collection (Burson, Actkinson, Hussey, & Jessup, 2012). A similar result was found in buffalograss, where multiple ploidy levels were found in the PI collections (Johnson et al., 1998).

The *F. ovina* complex includes seven species and three additional subspecies that are highly outcrossing and visually indistinguishable (Stace, 2010; Watson, 1958; Wilkinson & Stace, 1991). The 126 *F. ovina* USDA PI accessions used in our study have ploidy levels ranging from diploid to octoploid. The DNA content predicted by flow cytometry showed low average standard deviation, suggesting the methods for DNA content measurement is consistent. We measured DNA content variation between genotypes at same ploidy level, suggesting complex genome composition variation, which could be explained by post-polyploid diploidization events with differential gene loss (Mandáková & Lysak, 2018). This variation could also be the result of the change of transposon elements (Kidwell, 2002), and hybridization in the field (Leitch & Leitch, 2008; Soltis, Marchant, Van de Peer, & Soltis, 2015) or different chromosome size in different populations (Ceccarelli, Falistocco, & Cionini, 1992).

Although the DNA content of diploid samples was clearly separated from higher ploidies, there is no clear separation between accessions with higher ploidy levels. We also noticed ploidy level variation within some accessions. For example, of the four genotypes within the accession PI 330706 examined in this study, three were tetraploid while one was triploid. The triploid plant is likely the result of hybridization between the tetraploid plant with the pollen from some diploid relative. Another example is PI 235072, where we found three ploidy levels represented in the five genotypes we examined. It is known that *F. ovina* species can easily hybridize with relatives even at different ploidy levels (Jenkin & Jenkin, 1955). Under open pollination conditions, it is not surprising that the seed purity is low. Our results suggest that resources should be allocated such that the relevant USDA NPGS center managing open-pollinated species can properly isolate collections during seed increase.

While most accessions we surveyed can be assigned to discrete ploidy bins, 18% suggested DNA content that fell between ploidy levels. These accessions may have originated through hybridization between different diploid parents. It is also possible that these accessions are either aneuploid or dysploid, both of which have experienced chromosome gain/loss, or rearrangements. Further evaluation using a number of different approaches, ranging from morphological classification to genotyping, will be needed to fully classify these accessions.

In our study, all 16 accessions from China were found to be diploids and majority of plants from Iran were hexaploid. It is known that environmental and geographical factors played a role in *Festuca* ssp. genome size evolution (Ceccarelli et al., 1992; Šmarda, Bureš, Horová, Foggi, & Rossi, 2008). It would be interesting to see if the geographic location would be a factor to explain the distribution of ploidy levels. However, there is a lack of information on the specific area each wild accession was collected. Geographic information added to each accession in the NPGS collection would be useful; Rubenstein et al. (2006) found that NPGS users were more likely to utilize accessions when additional and accurate information was given about accessions. Beyond plant breeding, this new knowledge about the *F. ovina* collection might inspire other avenues of exploration such as investigating how geographic origin plays a role in taxon adaptation.

Seed size comparison suggests that accessions with higher ploidy levels tend to have a bigger seed size in *F. ovina* complex. Seed size often has an important impact on germination and plant development. Bretagnolle et al. (1995) found that larger seed size is correlated with higher ploidy level in *Dactylis glomerata* L. and the larger seed size had a positive influence on robust seedling growth (Bretagnolle, Thompson, & Lumaret, 1995). Larger seeds contain more carbohydrates that provide the seedling vigor to help increase the competitive advantage in the natural environment (Te Beest et al., 2011). Besides observing DNA content variation, we also observed phenotypic variation on leaf color within the *F. ovina* collection. Green and bluish leaf color was observed in all ploidy levels, suggesting that leaf-color-based fine fescue identification is not reliable (Beard, 1972; Hubbard, 1968).

## CONCLUSION

We evaluated 127 USDA PI accession and provided their DNA content and ploidy level estimation for 126 accessions. A total of 102 accessions were assigned to discrete ploidy levels, with the remaining had DNA content between discrete ploidy levels. Because of the cross-pollinating nature of *Festuca ovina* complex, better pollen control during the germplasm maintenance period could potentially reduce the chance of contaminations. Meanwhile, researchers should examine the PI collections to determine their ploidy level prior to adapting the accessions in their breeding program. This research builds the ground work for turfgrass researchers for using the *F. ovina* in their breeding program.

## Abbreviations

USDA: United States Department of Agriculture;
NPGS: National Plant Germplasm System;

## ACKNOWLEDGMENT

The authors would like to thank Dr. Ya Yang at the University of Minnesota for discussion about polyploid genome evolution. The authors would also like to thank Drs. Adrian Hegeman and Cory Hirsch at the University of Minnesota for reviewing this manuscript and providing comments and feedback. This research is funded by the National Institute of Food and Agriculture, U.S. Department of Agriculture, Specialty Crop Research Initiative under award number 2017-51181-27222.

## SUPPLEMENTAL MATERIAL

**Table S1.**
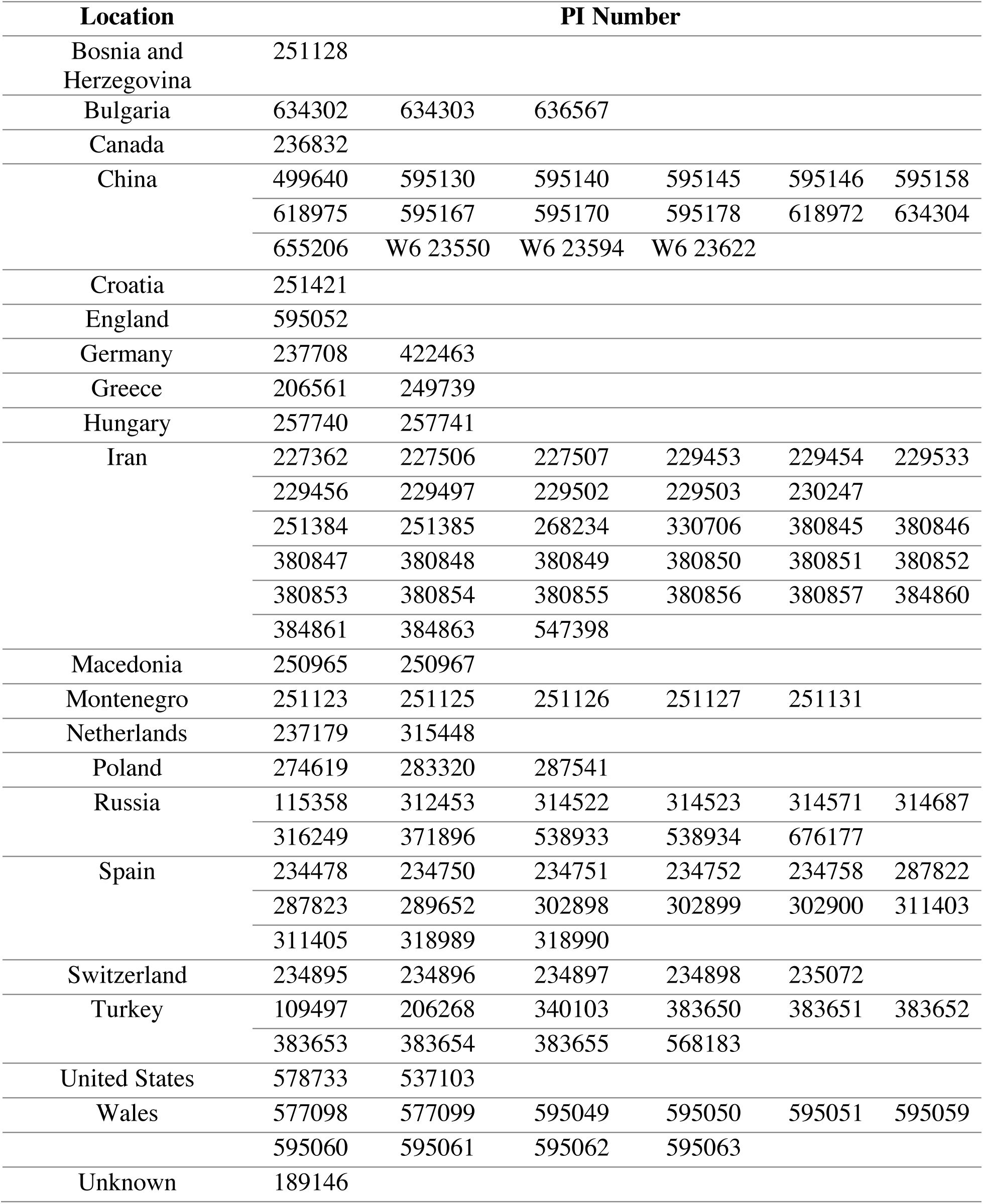
USDA PI collections by the country of origins. Accessions used in this study covered 20 countries, with Iran having the most entries.

| Location | PI Number |  |  |  |  |  |
| --- | --- | --- | --- | --- | --- | --- |
| Bosnia and Herzegovina | 251128 |  |  |  |  |  |
| Bulgaria | 634302 | 634303 | 636567 |  |  |  |
| Canada | 236832 |  |  |  |  |  |
| China | 499640 | 595130 | 595140 | 595145 | 595146 | 595158 |
|  | 618975 | 595167 | 595170 | 595178 | 618972 | 634304 |
|  | 655206 | W6 23550 | W6 23594 | W6 23622 |  |  |
| Croatia | 251421 |  |  |  |  |  |
| England | 595052 |  |  |  |  |  |
| Germany | 237708 | 422463 |  |  |  |  |
| Greece | 206561 | 249739 |  |  |  |  |
| Hungary | 257740 | 257741 |  |  |  |  |
| Iran | 227362 | 227506 | 227507 | 229453 | 229454 | 229533 |
|  | 229456 | 229497 | 229502 | 229503 | 230247 |  |
|  | 251384 | 251385 | 268234 | 330706 | 380845 | 380846 |
|  | 380847 | 380848 | 380849 | 380850 | 380851 | 380852 |
|  | 380853 | 380854 | 380855 | 380856 | 380857 | 384860 |
|  | 384861 | 384863 | 547398 |  |  |  |
| Macedonia | 250965 | 250967 |  |  |  |  |
| Montenegro | 251123 | 251125 | 251126 | 251127 | 251131 |  |
| Netherlands | 237179 | 315448 |  |  |  |  |
| Poland | 274619 | 283320 | 287541 |  |  |  |
| Russia | 115358 | 312453 | 314522 | 314523 | 314571 | 314687 |
|  | 316249 | 371896 | 538933 | 538934 | 676177 |  |
| Spain | 234478 | 234750 | 234751 | 234752 | 234758 | 287822 |
|  | 287823 | 289652 | 302898 | 302899 | 302900 | 311403 |
|  | 311405 | 318989 | 318990 |  |  |  |
| Switzerland | 234895 | 234896 | 234897 | 234898 | 235072 |  |
| Turkey | 109497 | 206268 | 340103 | 383650 | 383651 | 383652 |
|  | 383653 | 383654 | 383655 | 568183 |  |  |
| United States | 578733 | 537103 |  |  |  |  |
| Wales | 577098 | 577099 | 595049 | 595050 | 595051 | 595059 |

|  |  |  |  |  |
| --- | --- | --- | --- | --- |
|  | 595060 | 595061 | 595062 | 595063 |
| Unknown | 189146 |  |  |  |

**Table S2.** DNA content estimation and standard deviation of the USDA PI accessions. Data was sorted by the DNA content from the smallest to the largest.

| PI number | Average DNA Content (pg) | DNA Content SD (pg) | Estimated ploidy level |
| --- | --- | --- | --- |
| 109497 | 4.57 | 0.39 | 2 |
| 115358 | 3.77 | 0.19 | 2 |
| 189146 | 4.61 | 0.01 | 2 |
| 206268 | 4.51 | 0.40 | 2 |
| 206561 | 4.31 | 0.20 | 2 |
| 227362 | 9.77 | 1.04 | 4 |
| 227506 | 8.52 | 0.39 | 4 |
| 227507 | 8.49 | 0.27 | 4 |
| 229453 | 9.54 | 0.89 | 4 |
| 229454 | 8.39 | 0.37 | 4 |
| 229456 | 14.18 | 0.46 | 6 |
| 229497 | 8.72 | 0.35 | 4 |
| 229502 | 8.80 | 1.03 | 4 |
| 229503 | 14.45 | 0.64 | 6 |
| 229533 | 13.02 | 0.61 | 6 |
| 230247 | 13.26 | 0.48 | 6 |
| 234478 | 4.58 | 0.33 | 2 |
| 234750 | 18.07 | 0.36 | 8 |
| 234751 | 12.51 | 0.85 | 5 (5.32) |
| 234752 | 8.80 | 0.70 | 4 |
| 234758 | 9.74 | 0.61 | 4 |
| 234895 | 11.52 | 0.31 | 5 (4.90) |
| 234896 | 14.02 | 0.44 | 6 |
| 234897 | 4.20 | 0.26 | 2 |
| 234898 | 15.42 | 0.31 | 7 (6.56) |
| 236832 | 13.06 | 0.31 | 6 |
| 237179 | 4.22 | 0.24 | 2 |
| 237708 | 9.14 | 0.28 | 4 |
| 249739 | 4.78 | 0.18 | 2 |
| 250965 | 3.94 | 0.23 | 2 |
| 250967 | 13.41 | 0.09 | 6 |
| 251123 | 9.32 | 0.18 | 4 |
| 251125 | 10.18 | 0.92 | 4 |
| 251126 | 12.15 | 0.49 | 5 (5.17) |
| 251127 | 10.00 | 0.71 | 4 |
| 251128 | 9.14 | 0.27 | 4 |
| 251131 | 10.12 | 0.75 | 4 |
| 251384 | 4.97 | 0.15 | 2 |
| 251385 | 13.96 | 0.86 | 6 |
| 251421 | 4.74 | 0.19 | 2 |
| 257740 | 4.20 | 0.21 | 2 |
| 257741 | 4.64 | 0.15 | 2 |
| 268234 | 14.01 | 1.13 | 6 |
| 274619 | 12.86 | 1.25 | 5 (5.47) |
| 283320 | 10.14 | 0.29 | 4 |
| 287541 | 10.56 | 0.46 | 4 |
| 287822 | 9.42 | 0.92 | 4 |
| 287823 | 9.36 | 0.57 | 4 |
| 289652 | 7.67 | 0.20 | 3 (3.27) |
| 302898 | 9.31 | 0.32 | 4 |
| 302899 | 19.66 | 0.24 | 8 |
| 302900 | 9.03 | 0.33 | 4 |
| 311403 | 18.47 | 1.18 | 8 |
| 311405 | 12.61 | 1.20 | 5 (5.36) |
| 312453 | 10.49 | 0.71 | 4 |
| 314522 | 12.75 | 0.74 | 5 (5.43) |
| 314523 | 12.71 | 0.92 | 5 (5.41) |
| 314571 | 4.28 | 0.15 | 2 |
| 314687 | 4.26 | 0.32 | 2 |
| 315448 | 11.98 | 0.33 | 5 (5.10) |
| 316249 | 8.96 | 0.74 | 4 |
| 318989 | 19.43 | 0.76 | 8 |
| 318990 | 11.05 | 0.67 | 5 (4.70) |
| 330706 | 9.43 | 0.54 | 4 |
| 340103 | 9.26 | 0.27 | 4 |
| 371896 | 12.57 | 0.60 | 5 (5.35) |
| 380845 | 13.63 | 0.23 | 6 |
| 380846 | 13.25 | 0.30 | 6 |
| 380847 | 12.24 | 0.89 | 5 (5.21) |
| 380848 | 13.13 | 0.29 | 6 |
| 380849 | 15.39 | 0.37 | 7 (6.55) |
| 380850 | 12.31 | 0.62 | 5 (5.24) |
| 380851 | 13.79 | 0.33 | 6 |
| 380852 | 14.05 | 0.80 | 6 |
| 380853 | 14.35 | 0.62 | 6 |
| 380854 | 13.81 | 0.86 | 6 |
| 380855 | 14.11 | 1.12 | 6 |
| 380856 | 4.77 | 0.21 | 2 |
| 380857 | 13.83 | 0.24 | 6 |
| 383650 | 9.46 | 0.36 | 4 |
| 383651 | 4.87 | 0.10 | 2 |
| 383652 | 4.47 | 0.38 | 2 |
| 383653 | 8.15 | 0.23 | 3 (3.47) |
| 383654 | 8.38 | 0.65 | 4 |
| 383655 | 7.85 | 0.33 | 3 (3.34) |
| 384860 | 11.40 | 0.60 | 5 (4.85) |
| 384861 | 13.08 | 0.55 | 6 |
| 384863 | 14.98 | 0.25 | 6 |
| 422463 | 4.21 | 0.36 | 2 |
| 499640 | 4.35 | 0.18 | 2 |
| 537103 | 13.17 | 1.17 | 6 |
| 538933 | 11.69 | 0.32 | 5 (4.98) |
| 538934 | 4.17 | 0.34 | 2 |
| 547398 | 13.49 | 0.19 | 6 |
| 568183 | 4.22 | 0.23 | 2 |
| 577098 | 7.62 | 0.45 | 3 (3.24) |
| 577099 | 7.84 | 0.84 | 3 (3.33) |
| 578733 | 4.79 | 0.21 | 2 |
| 595049 | 7.62 | 0.41 | 3 (3.24) |
| 595050 | 9.27 | 0.48 | 4 |
| 595051 | 9.85 | 0.30 | 4 |
| 595052 | 9.34 | 0.50 | 4 |
| 595059 | 7.58 | 0.95 | 3 (3.23) |
| 595060 | 8.41 | 0.51 | 4 |
| 595061 | 9.36 | 0.74 | 4 |
| 595062 | 8.71 | 1.02 | 4 |
| 595063 | 8.52 | 0.43 | 4 |
| 595130 | 4.10 | 0.43 | 2 |
| 595140 | 4.30 | 0.30 | 2 |
| 595145 | 4.05 | 0.21 | 2 |
| 595146 | 3.82 | 0.04 | 2 |
| 595158 | 4.23 | 0.29 | 2 |
| 595167 | 4.25 | 0.04 | 2 |
| 595170 | 4.47 | 0.03 | 2 |
| 595178 | 4.34 | 0.29 | 2 |
| 618972 | 5.26 | 0.24 | 2 |
| 618975 | 4.08 | 0.12 | 2 |
| 634302 | 8.29 | 0.30 | 4 |
| 634303 | 4.47 | 0.17 | 2 |
| 634304 | 4.51 | 0.27 | 2 |
| 636567 | 3.98 | 0.28 | 2 |
| 655206 | 5.33 | 1.00 | 2 |
| 676177 | 4.63 | 0.32 | 2 |
| W6 23550 | 5.71 | 0.27 | 2 |
| W6 23594 | 5.63 | 0.09 | 2 |
| W6 23622 | 5.40 | 0.01 | 2 |

**Table S3.** The linear regression to associate seed size with ploidy levels. *t* statistics suggested there was a significant difference in seed size between different ploidy levels.

| Estimate | Std. | Error | *t* value | Pr (>\|*t*\|) |
| --- | --- | --- | --- | --- |
| **(Intercept)** | 2.1199 | 0.1079 | 19.644 | < 2.00E-16 *** |
| **Ploidy 4X** | 0.7584 | 0.1428 | 5.312 | 4.31E-07 *** |
| **Ploidy 6X** | 2.1432 | 0.1428 | 15.013 | < 2.00E-16 *** |
| **Ploidy 8X** | 2.3051 | 0.1526 | 15.104 | < 2.00E-16 *** |

Signif. codes: 0 '\*\*\*', 0.001 '\*\*', 0.01 '\*', 0.05 '.', 0.1 '.', 1

**Table S4.** The ANOVA analysis of PI accessions using ploidy levels as the variable. Significant differences were found between seed size and the corresponding ploidy level.

| Df | Sum | Sq | Mean | Sq | F- value | Pr(>F) |
| --- | --- | --- | --- | --- | --- | --- |
| **Ploidy** | 3 | 121.102 | 40.367 | 1.16E+02 | < 2.20E-16 | *** |
| **Residuals** | 136 | 47.514 | 0.349 |  |  |  |

Signif. codes: 0 '\*\*\*', 0.001 '\*\*', 0.01 '\*', 0.05 '.', 0.1 '.', 1

## AUTHOR CONTRIBUTIONS

YQ and SH performed the flow cytometry experiments, analyzed the data. JO performed the image analysis. YQ wrote the manuscript. EW secured funding for this project, supervised this research, provided suggestions, and comments. All authors contributed to the revision of the manuscript and approved the final version.

